# Myalgic Encephalomyelitis/Chronic Fatigue Syndrome and fibromyalgia are indistinguishable by their cerebrospinal fluid proteomes

**DOI:** 10.1101/2022.09.14.506792

**Authors:** Steven E. Schutzer, Tao Liu, Chia-Feng Tsai, Vladislav A. Petyuk, Athena A. Schepmoes, Yi-Ting Wang, Karl K. Weitz, Jonas Bergquist, Richard D. Smith, Benjamin H Natelson

## Abstract

Myalgic Encephalomyelitis/Chronic Fatigue Syndrome (ME/CFS) and fibromyalgia have overlapping neurologic symptoms particularly disabling fatigue. This has given rise to the question whether they are distinct central nervous system (CNS) entities or is one an extension of the other. To investigate this, we used unbiased quantitative mass spectrometry-based proteomics to examine the most proximal fluid to the brain, cerebrospinal fluid (CSF). This was to ascertain if the proteome profile of one was the same or different from the other. We examined two separate groups of ME/CFS, one with (n=15) and one without (n=15) fibromyalgia. We quantified a total of 2,083 proteins using immunoaffinity depletion, tandem mass tag isobaric labeling and offline two-dimensional liquid chromatography coupled to tandem mass spectrometry, including 1,789 that were quantified in all the CSF samples. ANOVA analysis did not yield any proteins with an adjusted p-value < 0.05. This supports the notion that ME/CFS and fibromyalgia as currently defined are not distinct entities.

## Introduction

Myalgic Encephalomyelitis/Chronic Fatigue Syndrome (ME/CFS) is an illness characterized by disabling fatigue, and fibromyalgia (FM) an illness characterized by body-wide pain. These two medically unexplained illnesses often exist together. This overlap has led some to consider the two illnesses to be part of the same illness spectrum^1^.

In contrast to this position, our own past data and studies from others found differences between the two illnesses^2,3^. That raised the possibility of different pathophysiological processes. These contrasting views compelled us to consider a study with newer unbiased technology.

Because our studies were subsumed under the auspices of an NIH-funded CFS Cooperative Research Center, we studied patients with ME/CFS only or with ME/CFS+FM.

We have viewed determining which of these hypotheses (same or distinct) is most likely as a testable research question with clinical impact. The question itself is clinically important. The existence of different pathophysiological processes will mean different paths to treatment. To investigate whether the two illnesses are the same or different, we used an approach that had helped us in an earlier study deconvolute the conundrum of whether ME/CFS was distinct from persistent neurologic Lyme disease syndrome (nPTLS) – i.e., mass spectrometry (MS)-based proteomics. Use of this technology has become the method of choice and discovery tool, to rapidly uncover protein biomarkers that can distinguish one disease from another^4-6^. The major features of both ME/CFS and nPTLS, just like ME/CFS and FM, are neurologic. We used MS-based proteomics to compare the cerebrospinal fluid (CSF) proteomes of both ME/CFS and nPTLS and were able to demonstrate that both were neurologic entities and distinct from one another and normals. We used the same approach here to compare ME/CFS with and without FM.

## Material and Methods

### Clinical specimens for this study

A total of 15 and 15 subjects were included in the CFS-only and CFS+FM groups, respectively. They were diagnosed as reported^7^. Ages were similar at 41.3 ± 9.4 SD and 40.1 ± 11.0, respectively. There were no differences in gender or rates of current comorbid psychiatric diagnosis between the groups. Samples were immediately aliquoted, frozen and stored at −80 ^o^C.

### Sample preparation and quantitative mass spectrometry methods

Two milliliters of each CSF sample was individually immunoaffinity depleted using an Agilent MARS Hu-14 column. The depleted samples were then reduced, alkylated, and tryptically digested using a trifluoroethanol-based single-tube approach without further sample cleanup. The resulting peptides were labeled using 11-plexed TMT reagents. A reference sample equally pooled from all 30 study samples were included with ten study samples in each of the three TMT-11 experiment. The TMT-labeled sample was fractionated using the high-pH reversed phase liquid chromatography (LC) and concatenated into 24 fractions. Each resulting fraction was analyzed using liquid chromatography coupled to tandem mass spectrometry (LC-MS/MS) on a Thermo Scientific Orbitrap Fusion Lumos Tribrid mass spectrometer. The LC-MS/MS data were processed using MSGF+ for identification of the peptides/proteins with a stringent cut of 1% FDR at both peptide and protein levels^8,9^. The TMT reporter ion intensity were extract using MASIC and used for protein quantification. The sample-to-reference ratios in each TMT experiment were log2 transformed, corrected for loading differences using medium centering, and batch corrected using the limma package in R. All statistical analyses, including ANOVA, unsupervised hierarchical clustering, and Random Forest analysis, were also performed in R using corresponding R packages.

## Results

Beginning with an unbiased proteomic approach where one does not need to know in advance what proteins may be in the sample, we performed comprehensive, quantitative proteomics using immunoaffinity depletion, tandem mass tag (TMT) isobaric labeling, and offline twodimensional liquid chromatography coupled to tandem mass spectrometry (LC-MS/MS)

**PCA analysis** of 1,789 proteins were quantified in all 15 ME/CFS-only (note labeled as CFS only in **Figure 1**) and 15 ME/CFS+FM (labeled CFS+FM in **Figure 1**) CSF samples. ANOVA analysis revealed a total of 14 proteins with p-value <0.05; however, none of these proteins has an adjusted p-value <0.05. Thus, there is no clear separation of the two groups using their entire CSF proteome profiles.

**Figure 1.**
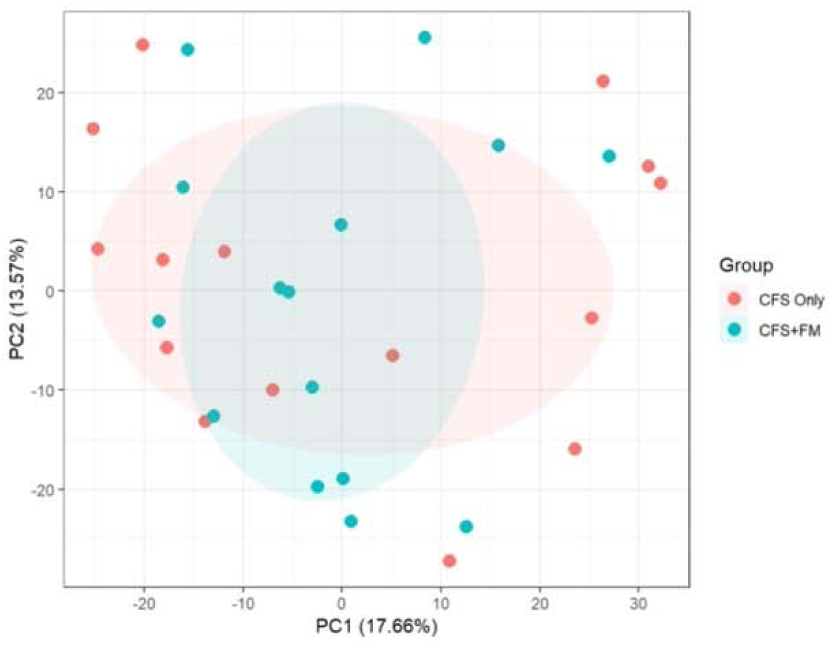
PCA analysis of 1,789 proteins quantified in all 15 CFS-only and 15 CFS+FM CSF samples. ANOVA analysis revealed a total of 14 proteins with p-value <0.05; however, none of these proteins has an adjusted p-value <0.05. There is no clear separation of the two groups using their entire CSF proteome profiles.

A Random Forest machine learning approach was then used to classify between the two groups based on relative protein abundance changes. Prediction accuracy was estimated using leave-one-out cross-validation (LOOCV). This did not distinguish the two groups either (**Figure 2**). The selection frequency for the differential proteins being selected into the Random Forest classifiers for separating the two groups is shown in the left panel (only 4 of 29 proteins were selected in 50% of the models), and the AUC composed from all LOOCV predictions was a modest 0.67 (right panel).

**Figure 2.**
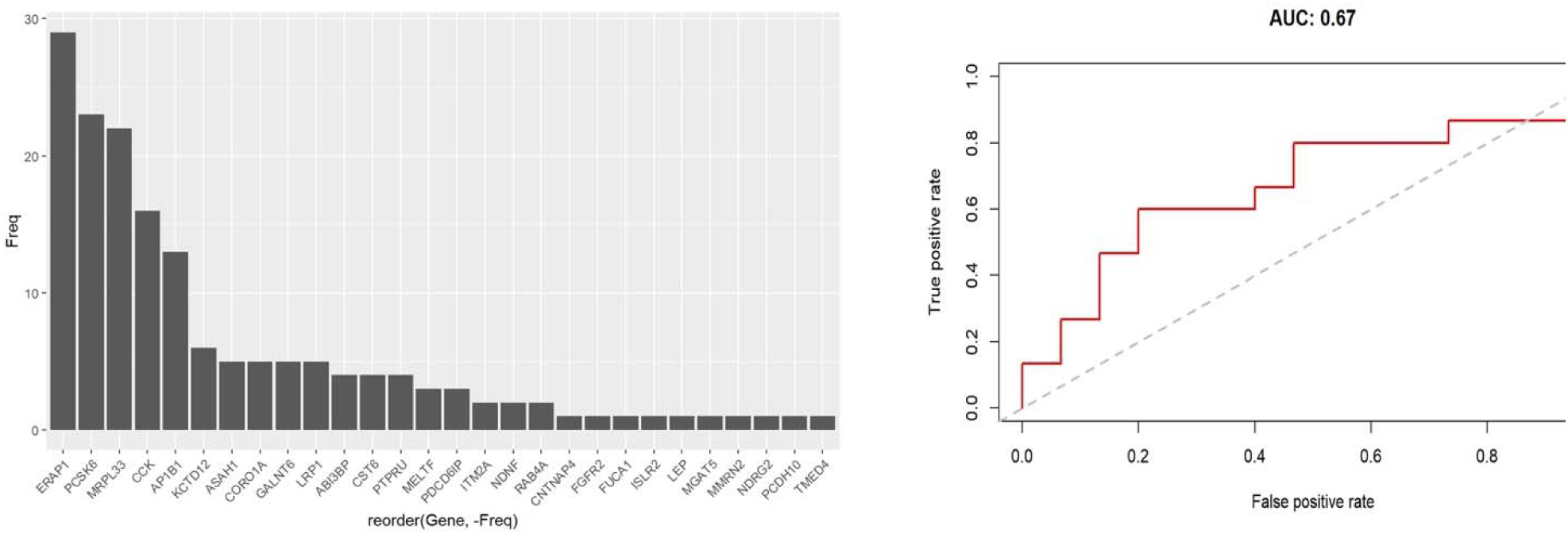
Random Forest classifier cross-validation. A Random Forest machine learning approach was used to classify between the two groups based on relative protein abundance changes. Prediction accuracy was estimated using leave-one-out cross-validation (LOOCV). The selection frequency for th differential proteins being selected into the Random Forest classifiers for separating the two groups is shown in the left panel (only 4 of 29 proteins were selected in 50% of the models), and the AUC composed from all LOOCV predictions was 0.67 (right panel).

Lastly, we employed unsupervised hierarchical clustering analysis, and there is no clear separation of the two groups using their entire CSF proteome profiles (**Figure 3**).

**Figure 3.**
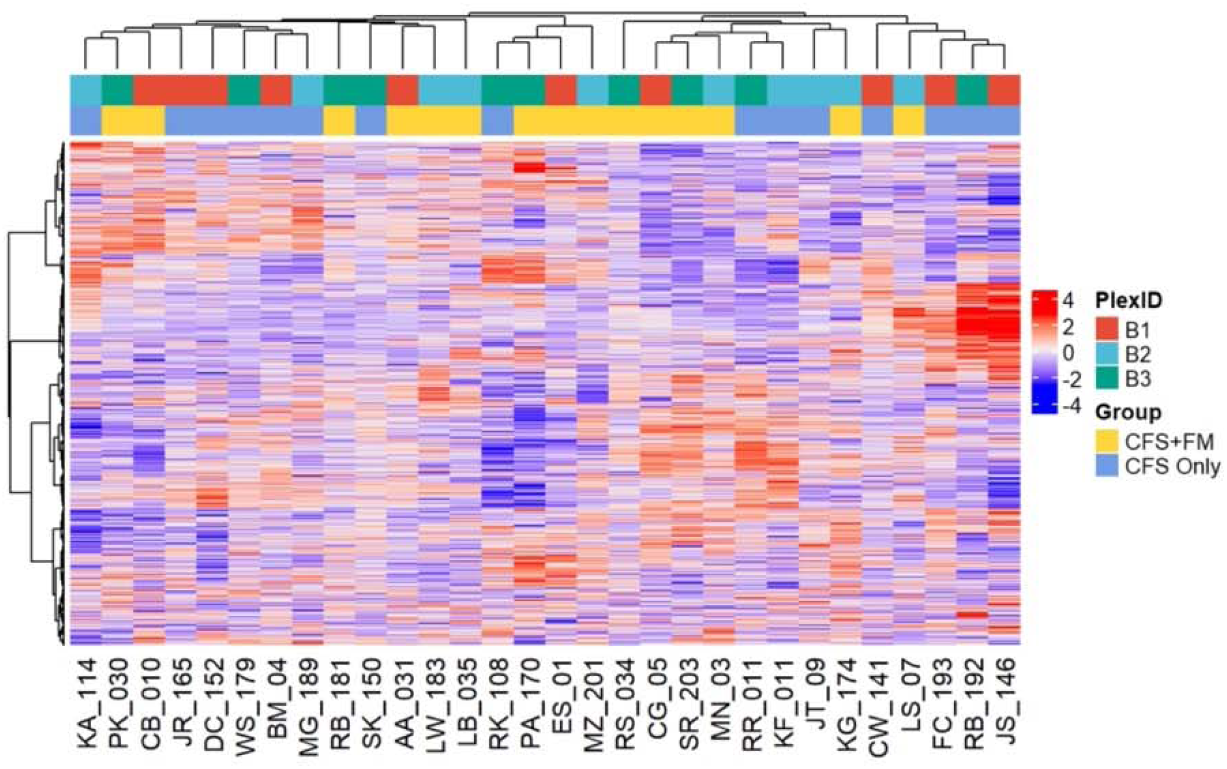
Unsupervised hierarchical clustering of the entire CSF proteome. There is no clear separation of the two groups using their entire CSF proteome profiles. The TMT-11 plex from which the individual samples were analyzed is also shown as a reference.

Unsupervised hierarchical clustering of the top 20% most variable proteins in CSF proteome did not lead to clear separation of the two groups either (**Figure S1**).

The sum of these results does not support the hypothesis that ME/CFS and ME/CFS with fibromyalgia are distinct entities. Even with comprehensive proteome coverage and precise quantification, no distinguishing CSF proteins could be found between patients with ME/CFS and those with both ME/CFS + FM.

## Discussion

CFS, now known as ME/CFS, and FM are medically unexplained illnesses, predominantly of women, characterized by disabling fatigue and by widespread pain with tenderness, respectively. The prevalence of ME/CFS is a tenth that of FM (0.3%^10^ vs 3%^11^) because, in contrast to FM which has no exclusions, ME/CFS is not diagnosed in the face of any disease process that could produce fatigue. However, the core symptoms of pain, fatigue, sleep problems and cognitive difficulties exist across both syndromes and lead to significant comorbidity between them. The fact that these two syndromes co-exist so often has led some to question whether they are, in fact, distinct diagnostic entities. Wessely et al.^1^ suggested that the “similarities between them outweigh the differences” – a position taken by other researchers^12^. As noted by Hauser, FM is rarely a stand-alone condition. A revision of the original 1990 case definition for FM published in 2010^13^ has reduced the difference between the two illnesses with nearly twice as many ME/CFS patients also receiving the diagnosis of FM than when the 1990 case definition is used^14^. Blurring the difference between the two syndromes left open the research question of whether the two illnesses are spectrum variants of one another or are due to distinct different pathophysiological processes which this current study examined.

The quantitative proteomics approach we used examining the CSF as a reflection of the CNS-related major symptoms did not support the hypothesis that the two conditions studied – ME/CFS with and without fibromyalgia – were separable. Thus, there do not appear to be discrete CSF proteins for ME/CFS which allow that group of patients to be clearly differentiated from those for ME/CFS+FM.

We and others have had success in distinguishing one disease from another beginning with the first step: an unbiased discovery using MS-based proteomics that employs immunoaffinity depletion of common abundant proteins that could otherwise mask the less abundant proteins that have higher biomarker potential^7,15^, followed by isobaric labeling of the resulting low abundance proteins in each individual sample. In the current study the same methods that potentially could have distinguished one disease from another, did not demonstrate a pathophysiologic distinction between ME/CFS and ME/CFS+FM. Although we did not have the benefit of the rare resource of CSF from “pure” FM group, we expected differences to show up by skewing from the ME/CFS+FM group. As shown in **Figures 1, 2, 3, and S1** no statistically significant differences were found. This basic science analysis comparing the two illness processes suggests that the two may share a similar pathophysiological basis. If that conclusion is supported by other research including a set of FM only patients, this would support the notion that the two illnesses fall along a common illness spectrum and may be approached as a single entity – with implications for both diagnosis and the development of new treatment approaches.

## Acknowledgments

Funding was provided by a grant from National Institutes of Health (NIH) and National Institute of Allergy and Infectious Disease (NIAID) AI135879.

## Author Contributions

SES, TL, BHN, RDS, JB. contributed to the conception and design of the study; SES, TL, CT, VAP, AAS, YW, KKW, RDS, BHN, contributed to the acquisition and analysis of data; SES, TL, CT, BHN contributed to drafting the text or preparing the figures.

---

**Figure S1.**
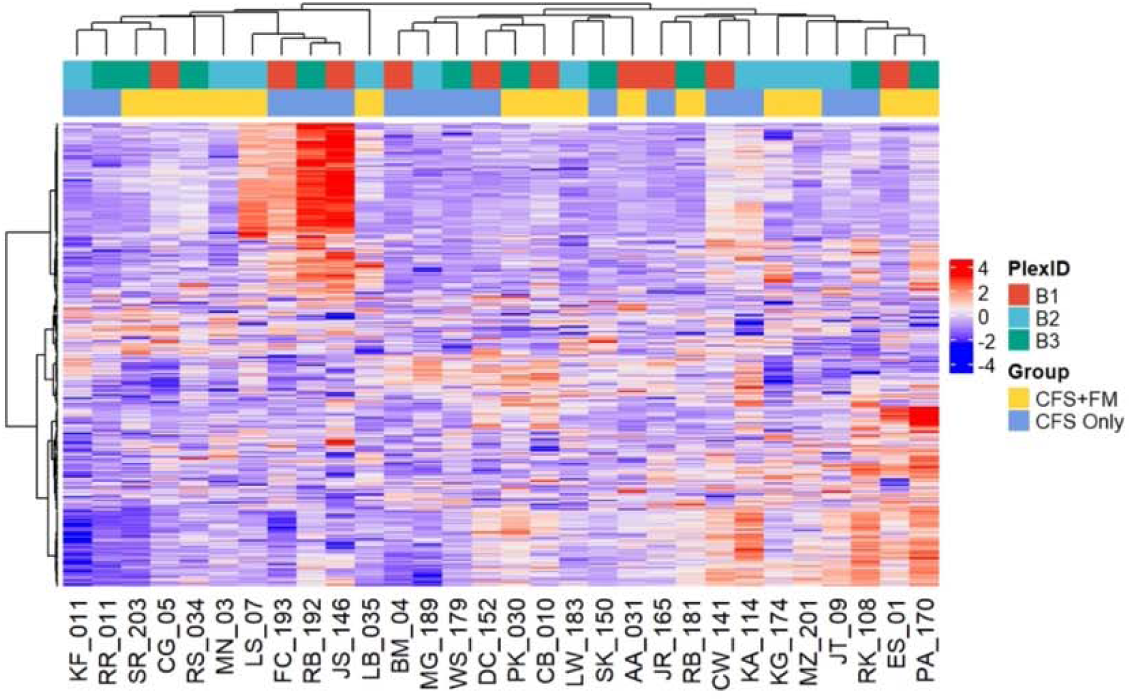
Unsupervised hierarchical clustering of the top 20% most variable CSF proteome. There is no clear separation of the two groups. TheTMT-11 plex from which the individual samples were analyzed is also shown as a reference.

